# The ABA INSENSITIVE (ABI) 4 transcription factor is stabilized by stress, ABA and phosphorylation

**DOI:** 10.1101/2022.02.01.478625

**Authors:** Tzofia Maymon, Nadav Eisner, Dudy Bar-Zvi

## Abstract

The Arabidopsis transcription factor ABSCISIC ACID INSENSITIVE 4 (ABI4) is a key player in the plant hormone abscisic acid (ABA) signaling pathway. ABI4 is also involved in seed development and germination, the response to abiotic stresses such as drought and salinity, control of lipid reserve mobilization in the embryo, lateral root formation, and redox control. Expression of the *ABI4* gene is tightly regulated and basal expression is low. Maximal transcript levels occur during seed maturation and in the early stages of seed germination and are markedly reduced in other developmental stages. ABI4 is an unstable lowly expressed protein, resulting from tight post-transcriptional regulation. Here, we studied factors affecting the stability of the ABI4 protein using transgenic Arabidopsis plants expressing *35S::HA-FLAG-ABI4-eGFP*. Despite the expression of eGFP-tagged ABI4 being driven by the highly active 35S CaMV promoter the steady-state levels of ABI4 were extremely low in the roots of seedling grown in optimal conditions. These levels were markedly enhanced upon exposure of the seedlings to abiotic stress and ABA. ABI4 is degraded rapidly by the 26S proteasome and we report on the role of phosphorylation of ABI4-serine 114 in regulating ABI4 stability. Our results indicate that ABI4 is tightly regulated both post-transcriptionally and post-translationally. Moreover, abiotic factors and plant hormones have similar effects on ABI4 transcripts and ABI4 protein levels. This double-check mechanism for controlling ABI4 reflects on its central role in plant development and cellular metabolism.

**SIGNIFICANCE STATEMENT:** We show that stabilization of the ABI4 transcription factor by stress and hormones is mediated by phosphorylation of Serine 114 by MAP kinases. Transcription of *ABI4* is also modulated by MAP kinases, suggesting that the same signals affect both transcript and protein levels, resulting in tight modulation of ABI4 activity.

## INTRODUCTION

Plant development and response to environmental cues involve signaling pathways in which the last components are often transcription factors (reviewed by (Chen *et al*., 2021) As a result, these signaling pathways affect the transcription of a large number of genes whose expression is affected by the respective transcription factors.

The Arabidopsis *ABSCISIC ACID INSENSITIVE 4 (ABI4)* gene encodes an Apetala 2 (AP2) family transcription factor (Finkelstein *et al*., 1998). Apetala 2 is a ∼60 amino acid long plant-specific DNA-binding domain, characterized first in the Arabidopsis *APETALA2* homeotic gene (Jofuku *et al*., 1994; Okamuro *et al*., 1997). The *ABI4* gene was identified by screening of gamma-irradiated Arabidopsis seeds for mutants capable of germination in the presence of inhibitory concentrations of the plant hormone abscisic acid (ABA) (Finkelstein, 1994). ABI4 alleles were isolated by screening for germination in the presence of high concentrations of salt and sugar (Arenas-Huertero *et al*., 2000; Huijser *et al*., 2000; Laby *et al*., 2000; Quesada *et al*., 2000; Rook *et al*., 2001). ABI4 also plays a central role in other plant signaling pathways, including lipid mobilization, lateral root development, regulation of light-modulated genes, redox signaling, pathogen response, and mitochondrial retrograde signaling (reviewed in Wind *et al*., 2013). Its role in chloroplast retrograde signaling is disputed (Koussevitzky *et al*., 2007; Kacprzak *et al*., 2019).

*ABI4* expression is tightly developmentally regulated: the highest steady-state levels of ABI4 transcript are in embryos, maturing pollen, and early germination stages (Söderman *et al*., 2000; Nakabayashi *et al*., 2005; Penfield *et al*., 2006). The transcript levels are significantly reduced in other developmental stages; its expression is restricted to root phloem companion cells and parenchyma, and to some extent to the vascular system of the shoot (Shkolnik-Inbar and Bar-Zvi, 2010; Shkolnik-Inbar and Bar-Zvi, 2011; Shkolnik-Inbar *et al*., 2013). In addition, steady-state levels of *ABI4* transcript levels are enhanced by ABA, NaCl, and glucose and repressed by auxin (Arroyo *et al*., 2003; Shkolnik-Inbar and Bar-Zvi, 2010; Shkolnik-Inbar *et al*., 2013).

ABI4 is a highly unstable protein (Finkelstein *et al*., 2011; Gregorio *et al*., 2014). Several protein motifs, such as PEST and AP2-associated (Finkelstein *et al*., 2011; Gregorio *et al*., 2014), destabilize ABI4 via degradation by the proteasome. Other regions of the protein destabilize it in a proteasome-independent manner (Finkelstein *et al*., 2011). ABI4 is stabilized by high concentrations of salt and sugar (Finkelstein *et al*., 2011; Gregorio *et al*., 2014), and by preventing light exposure (Xu *et al*., 2016). COP1 is involved in the light mediated degradation of ABI4 (Xu *et al*., 2016): levels of ABI4 were enhanced in light-exposed *cop1* mutant seedlings, and further increased by treating *cop1* mutants with the MG132 proteasome inhibitor, suggesting that COP1, as well as additional E3s, modulate ABI4 stability (Xu *et al*., 2016).

Being downstream of the signaling pathway cascades, transcription factors are frequently modulated by phosphorylation resulting in their activation or inhibition. ABI4 was phosphorylated *in vitro* by MPK3, MPK4, and MPK6 (Popescu *et al*., 2009; Guo *et al*., 2016; Bai *et al*., 2020; Eisner *et al*., 2021). Phosphorylation of ABI4 by MAPKs repressed the expression of the *LHCB* gene (Guo *et al*., 2016), and inhibited the emergence of adventitious roots (Bai *et al*., 2020). In addition, the phosphorylation of S114 is essential for the biological activity of ABI4 as shown in studies of the complementation of the *abi4* mutant phenotype (Eisner *et al*., 2021).

Here, we studied factors affecting the stability of the ABI4 protein in Arabidopsis plants by expressing *HA-FLAG-ABI4-eGFP* driven by the constitutive highly active 35S promoter. The tagged ABI4 could be detected in embryos rescued from imbibed seeds but not in seedlings. Treatment of the seedlings with NaCl resulted in a transient stabilization of ABI4 peaking at 2-4 h. ABA and high glucose also stabilized ABI4-eGFP but with slower kinetics and reached lower levels than NaCl treatment. The phosphomimetic ABI4 (S114E) protein was more stable than the wild type ABI4 and the phosphorylation-null ABI4 (S114A) mutant in salt-treated plants, suggesting that phosphorylation of ABI4 by MAPKs results in stabilization of ABI4. Interestingly, NaCl, ABA, and glucose are known to affect similarly the steady-state levels of ABI4 transcripts (Arroyo *et al*., 2003; Shkolnik-Inbar and Bar-Zvi, 2010; Shkolnik-Inbar *et al*., 2013). We thus propose that the MAPK signaling cascade also activates ABI4 transcription via the phosphorylation of MYB, WORKY, and ABI4 transcription factors known to transactivate the transcription of the *ABI4* gene. As a result, similar cues regulate *ABI4* on transcriptionally and posttranscriptionally levels, resulting in a very tight regulation of this key factor.

## RESULTS

### The *35S::HA-FLAG-ABI4-eGFP* construct encodes a biologically active protein

To study ABI4 *in planta* we used the enhanced Green Fluorescent Protein (eGFP) (Cinelli *et al*., 2000) fused to the carboxy terminus of ABI4, and transcription driven by the highly active cauliflower mosaic virus constitutive 35S promoter (35S) (Odell *et al*., 1985). We previously found that overexpressing *35S::ABI4* in Arabidopsis resulted in seedling death within three days of germination, whereas fusing the HA_3_-FLAG_3_ tag to the N-terminus of ABI4 resulted in viable plants (Shkolnik-Inbar and Bar-Zvi, 2010). We thus constructed *35S::HA-FLAG-ABI4-eGFP* and used it for the transformation of Arabidopsis. To determine if HA-FLAG-ABI4-eGFP protein is biologically active, we tested whether tagged ABI4 can complement the *abi4-1* mutant. *abi4-1* is a frameshift mutant resulting from a single bp deletion at codon 157 (Finkelstein *et al*., 1998); the expressed protein has the AP2 DNA binding domain but lacks the transactivation domain. Homozygous transgenic plants, resulting from a single T-DNA insertion in the genome of the *abi-4* mutants did not have any visible phenotype when grown on agar plates with 0.5 x MS, 0.5% sucrose medium, or in pots containing potting mix. To determine if the expressed HA-FLAG-ABI4-eGFP (ABI4-eGFP) protein retains the biological activity of ABI4, we examined its ability to complement the phenotype of *abi4* mutants by assaying seed germination in the presence of ABA, the best-studied phenotype of these mutants (Finkelstein, 1993). Figure 1 shows that expressing HA-FLAG-ABI4-eGFP in *abi4-1* plants restored the ABA sensitivity, indicating that tagging ABI4 at both its amino- and carboxy- termini does not impair its biological activity.

**Figure 1.**
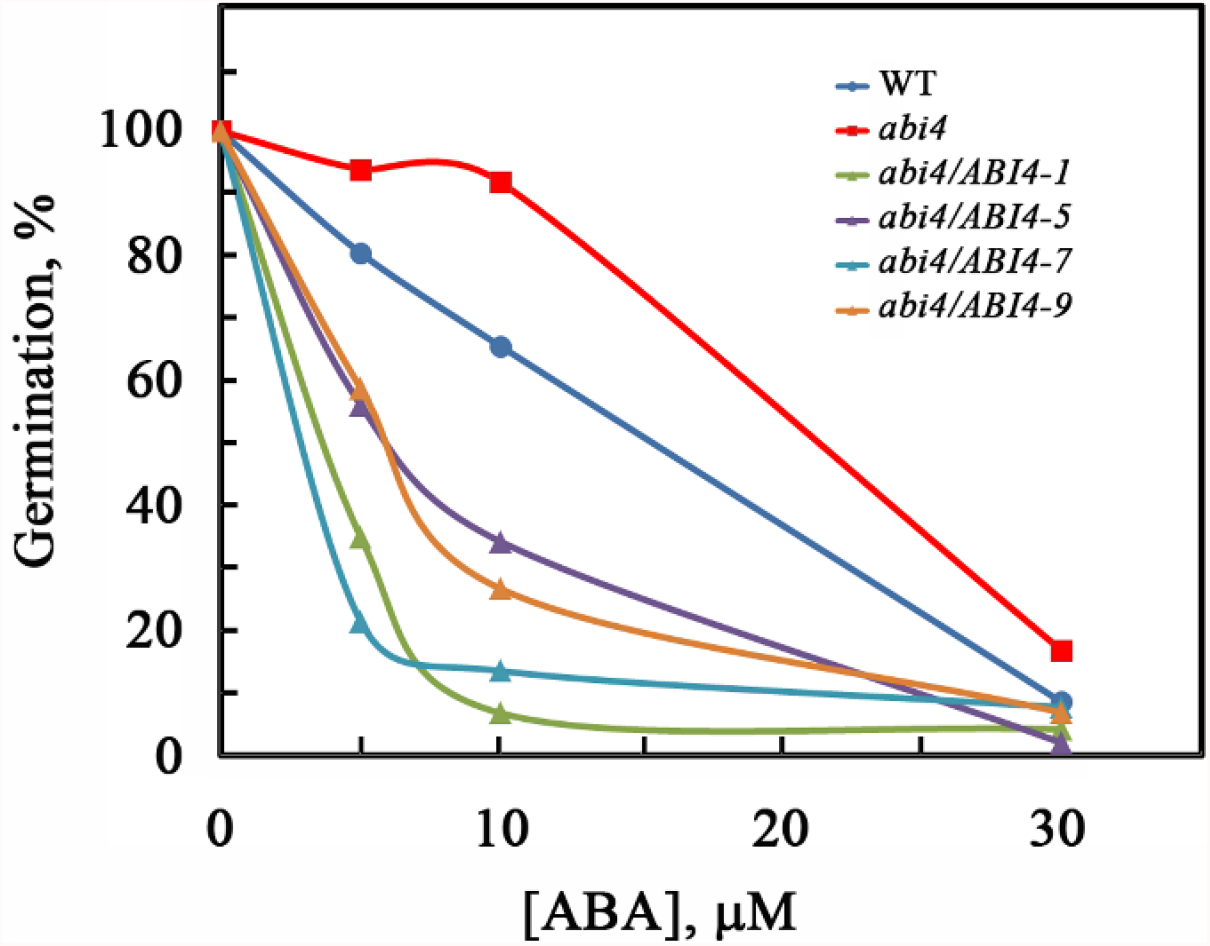
Complementation of the *abi4-1* mutant by *35S::HA*_*3*_*-FLAG*_*3*_*-ABI4-eGFP*. Seeds of the homozygous plants of the indicated genotypes were plated on agar-solidified 0.5 x MS, 0.5% sucrose medium supplemented with the indicated concentrations of ABA. Germination was scored 7-days later. *abi4, abi4-1* mutant; *abi4/ABI4-1, −5, −7, −9*, transgenic lines 1, 5, 7, 9 of *abi4-1* plants transformed with the *35S::HA*_*3*_*-FLAG*_*3*_*-ABI4-eGFP* construct.

The 35S promoter is a commonly used strong constitutive promoter that is active in most plant tissues (Amack and Antunes, 2020). We thus expected to detect high eGFP fluorescence signals in seedlings of WT plants transformed with the *35S::HA-FLAG-ABI4-eGFP* construct (WT/ABI4-eGFP). Surprisingly, we could not detect significant fluorescent signals in these plants (Figure 2A). To confirm the construct, we examined the fluorescence in embryos prepared from imbibed seeds, where we could detect a high fluorescent signal (Figure 2C). We interpret these results as an indication that ABI4 is subjected to post-transcriptional regulation and that ABI4 protein levels are tightly regulated.

**Figure 2.**
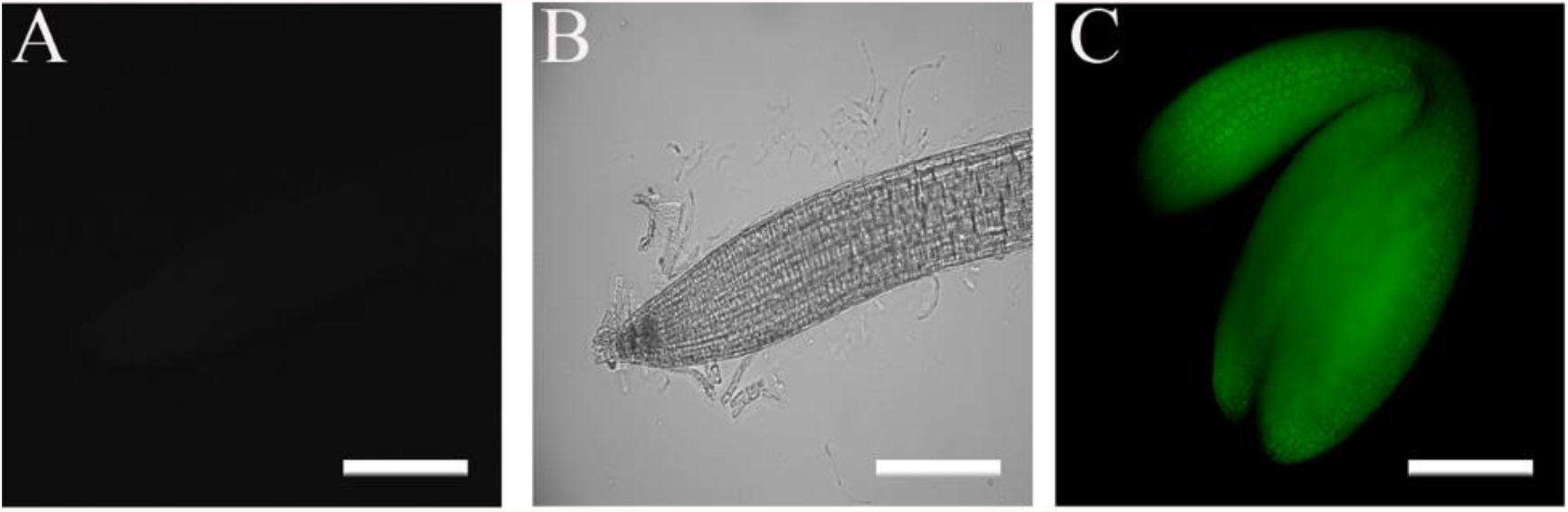
Verification of the transgenic lines. The fluorescence of plants transformed with the *35S::HA*_*3*_*-FLAG*_*3*_*-ABI4-eGFP* construct examined by fluorescent microscopy. (A, B) ten day old root: (A) fluorescence image, (B) a bright field image (C) fluorescence image of embryo extracted from 24-h imbibed seed.

### Accumulation of ABI4-eGFP is NaCl dependent

Previous studies showed that environmental signals post-transcriptionally regulate ABI4. To examine if different external and internal cues affect the steady-state levels of ABI4, we tested cues known to affect the activity of the *ABI4* promoter. The steady-state levels of *ABI4* mRNA driven by its endogenous promoter are enhanced by NaCl (Shkolnik-Inbar *et al*., 2013). We thus examined if NaCl also affects protein levels of ABI4 when transcription is driven by the 35S promoter. Exposing WT plants expressing *ABI4-eGFP* to 0.3 M NaCl resulted in a transient increase in the eGFP-fluorescence signal, with the maximal signal observed 2-3 h following seedling exposure to salt (Figure 3A). The signal was NaCl dose dependent with the maximum at 0.3 M NaCl (Figure 3B). No fluorescence was observed in seedlings transferred to fresh 0.5 x MS, 0.5% sucrose medium, suggesting that the transient increase in fluorescence seen in the NaCl treated seedlings did not result from transferring the seedlings from the agar plates to the buffer soaked filter paper. To confirm the observed fluorescence signals, protein extracts of roots of salt-treated WT/*ABI4-eGFP* seedlings were subjected to western blot analysis using an anti-GFP antibody. The results confirmed that ABI4-eGFP is essentially not detectable in control non-treated roots, whereas a transient increase of ABI4-eGFP can be seen following exposure to NaCl peaking at 2 h after the application of NaCl (Figure 3C). The protein levels of ABI4-eGFP were low, even at maximal values, and could be only detected with a high-sensitive detection assay.

**Figure 3.**
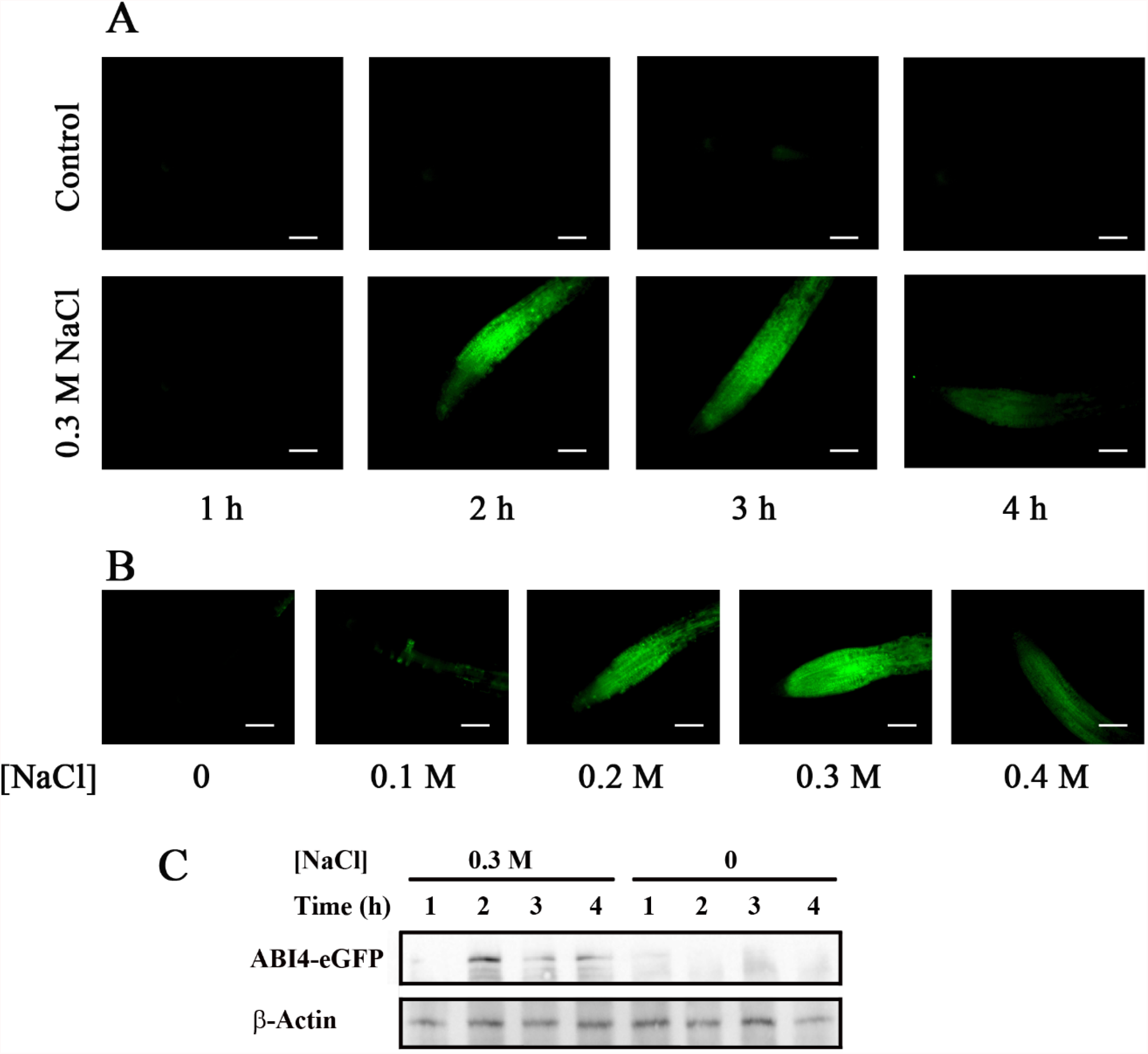
NaCl treatment transiently enhances ABI4-eGFP protein levels. Ten days old Arabidopsis plants expressing the *35S::HA*_*3*_*-FLAG*_*3*_*-ABI4-eGFP* construct incubated for the indicated times with 0.5 x MS salts, 0.5% sucrose without (A, control, upper row) or with 0.3 M NaCl (A, lower row), or for 2.5 h with growth medium containing the indicated incubated concentrations of NaCl (B) Roots were examined by fluorescence microscopy. Scale bar = 100 μm. (C) Western blot analysis showing the expression levels of ABI4-eGFP following NaCl treatment. β-actin used as loading control.

The NaCl-dependent increase in ABI4 protein levels may result from either changes in the transcript levels of the encoding mRNA or regulation of the protein levels. To assess this point, we quantified the *ABI4-eGFP* transcript and protein levels in roots of untreated and NaCl-treated seedlings. *ABI4-eGFP* transcript levels were determined by RT-qPCR using amplification primers from the sequence encoding the HA_3_-FLAG_3_ tag to avoid assaying the expression of the endogenous *ABI4* gene. Following treatment with 0.3 M NaCl, the steady-state mRNA levels of *ABI4-eGFP* were increased 2.0 and 1.6 fold at 2.5 and 4 h, respectively. ABI4-eGFP protein levels quantified by using the fluorescence intensity of the roots were 32.7 and 7.3 times higher for roots of plants exposed to 0.3 M NaCl for 2.5 and 4 h, respectively, compared to control untreated roots (Figure 4). Control fluorescence signals of plants expressing *35S::GFP* were not affected by salt treatment (Figure S1). These results indicate that the NaCl-dependent increase in ABI4-eGFP protein levels results from post-transcriptional regulation of *ABI4* rather than from changes in the transcript levels or the salt effect on the GFP tag.

**Figure 4.**
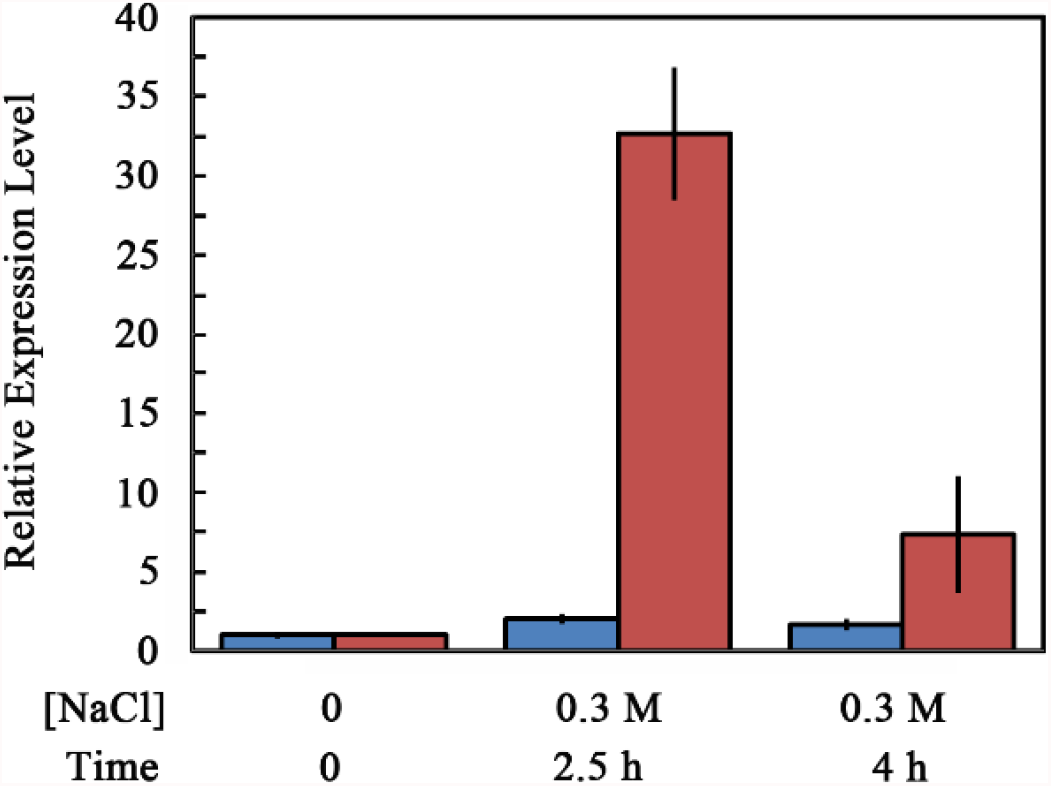
Effect of NaCl treatment on steady state levels of *ABI4-eGFP* transcript and protein in the roots. Ten days old seedlings transformed with the *35S::HA*_*3*_*- FLAG*_*3*_*-ABI4-eGFP* construct were transferred onto a filter paper soaked with 0.5 x MS salts, 0.5% sucrose, with or without 0.3 M NaCl. Roots were harvested at the indicated times, and the levels of *HA*_*3*_*-FLAG*_*3*_*-ABI4-eGFP* transcript (blue) or protein (red) were determined by RT-qPCR and fluorescence microscopy, respectively.

### Subcellular localization of ABI4-eGFP following NaCl treatment is cell-type-specific

Although NaCl treatment of plants expressing the *35S::HA-FLAG-ABI4-eGFP* construct resulted in enhanced protein levels in most root cells, the observed fluorescence pattern of the ABI4-eGFP was diffusive in most cell types. In contrast, it was found in spherical structures in root stele suggesting nuclear localization (Figure 5A). Staining the roots of NaCl-treated *ABI4-eGFP* plants with the DNA fluorescence stain 4′,6-diamidino-2-phenylindole (DAPI) shows that the DAPI and eGFP fluorescence signals overlap, confirming that ABI4-eGFP is localized in the nuclei of root stele cells (Figure 5C). This pattern is specific to ABI4 since it was not observed in the roots expressing the eGFP-tag alone (Figure S1).

**Figure 5.**
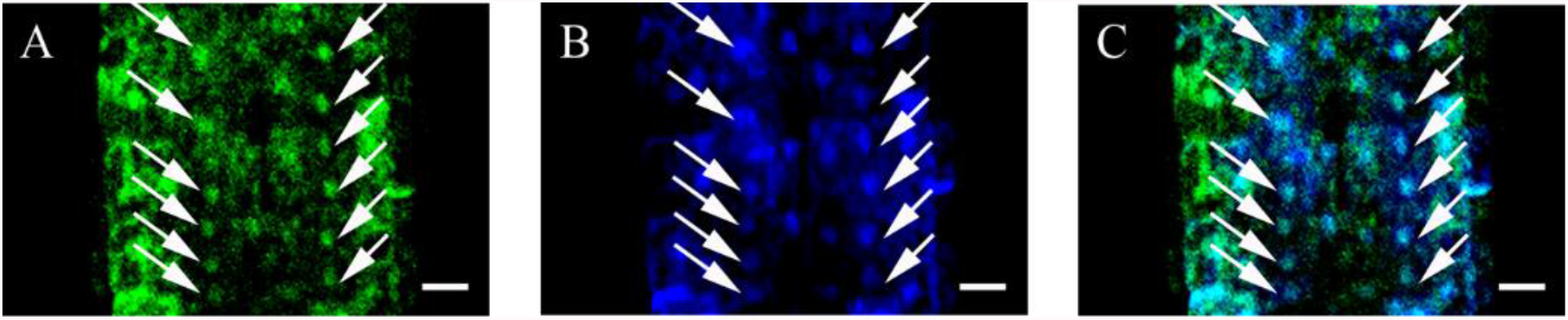
The subcellular localization of ABI4-eGFP in the roots is cell type dependent. Ten day old seedlings expressing the *35S::HA*_*3*_*-FLAG*_*3*_*-ABI4-eGFP* construct were treated with 0.3 M NaCl for 2.5 h. Roots were stained with DAPI and examined by confocal microscopy. (A) GFP fluorescence (B) DAPI fluorescence (C) merged image of A and B. Arrows point to nuclei. Scale bar = 10 μm.

### ABA and glucose treatment enhance ABI4-eGFP protein levels

Transcript levels of endogenous *ABI4* are also induced by treatment with ABA or high concentrations of glucose (Arroyo *et al*., 2003). To determine if these treatments also affect the levels of the ABI4-eGFP protein, ten day old *35S::HA-FLAG-ABI4-eGFP* transgenic plants were transferred to media containing ABA or glucose, and ABI4-eGFP accumulation was followed by fluorescence microscopy. Enhanced ABI4-eGFP levels were detected in the root stele of seedlings treated with ABA or glucose approximately 24 after treatments (Figure 6).

**Figure 6.**
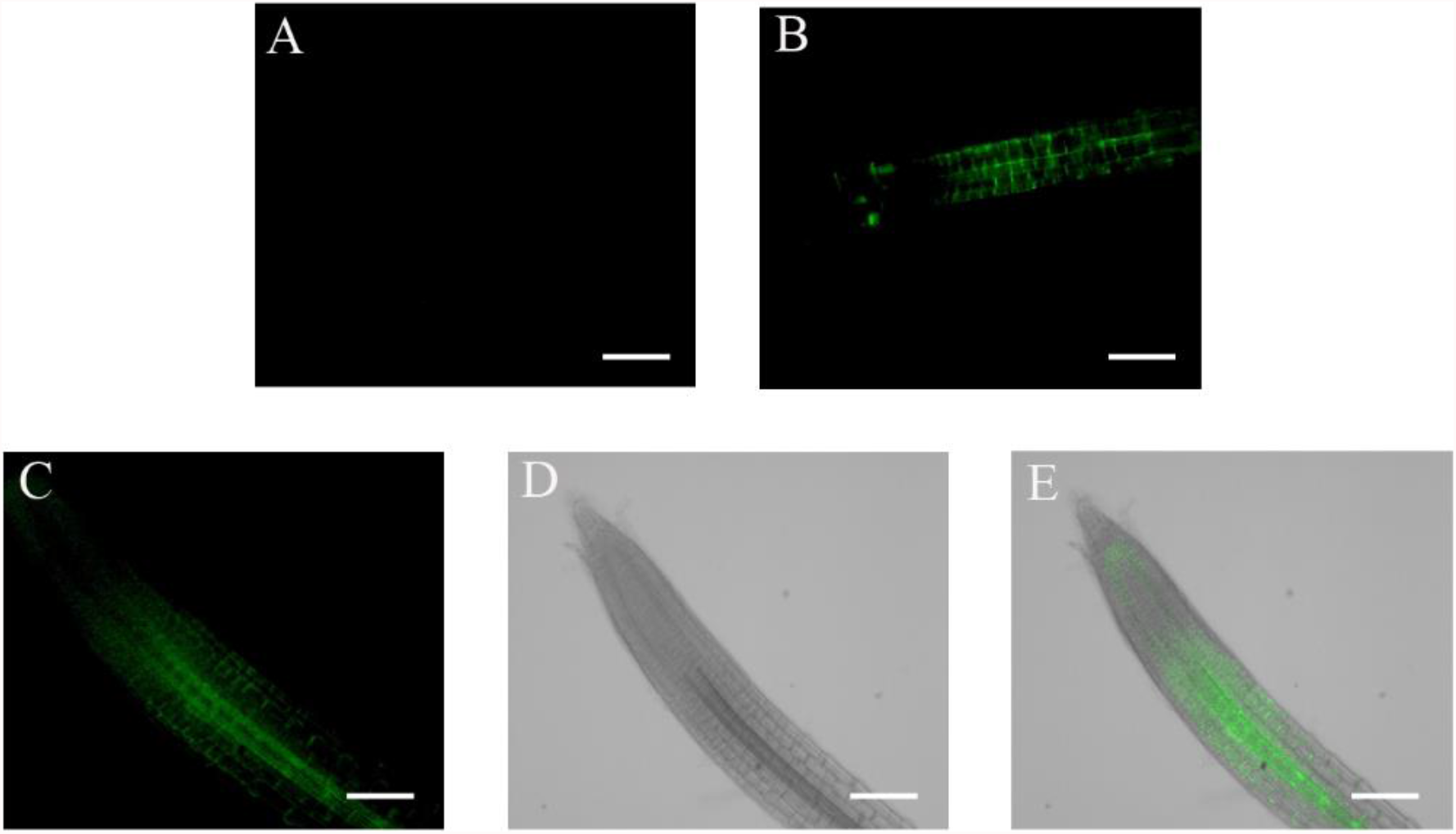
ABI4 expression is increased following glucose and ABA treatment. Ten days old *35S::HA*_*3*_*-FLAG*_*3*_*-ABI4-eGFP* expressing plants were incubated for 24 h in 0.5 x MS, 0.5% sucrose growth medium (A), or in the same medium supplemented with 7% Glucose (B) or 30 μM ABA (C). (D and E) Bright-field and the merged image of the ABA treated root shown in (C). Scale bar = 100 μm.

### Auxin counteracts the NaCl- induced increase of ABI4-eGFP levels

ABI4 mediates cytokinin inhibition of lateral roots formation by reducing the polar transport of auxin, a plant hormone known to induce the formation of lateral roots (Shkolnik-Inbar and Bar-Zvi, 2010). Exogenous auxin also reduced the steady-state levels of *ABI4* transcripts in the roots (Shkolnik-Inbar and Bar-Zvi, 2010). To test if auxin also post-transcriptionally regulates ABI4, we tested if auxin counteracts the NaCl-induced enhancement of ABI4-eGFP. Figure 7 shows that when added together with NaCl, 3-indole acetic acid (IAA) prevented the NaCl-induced accumulation of ABI4-eGFP, indicating that auxin negatively regulates the steady-state levels of the ABI4 protein.

**Figure 7.**
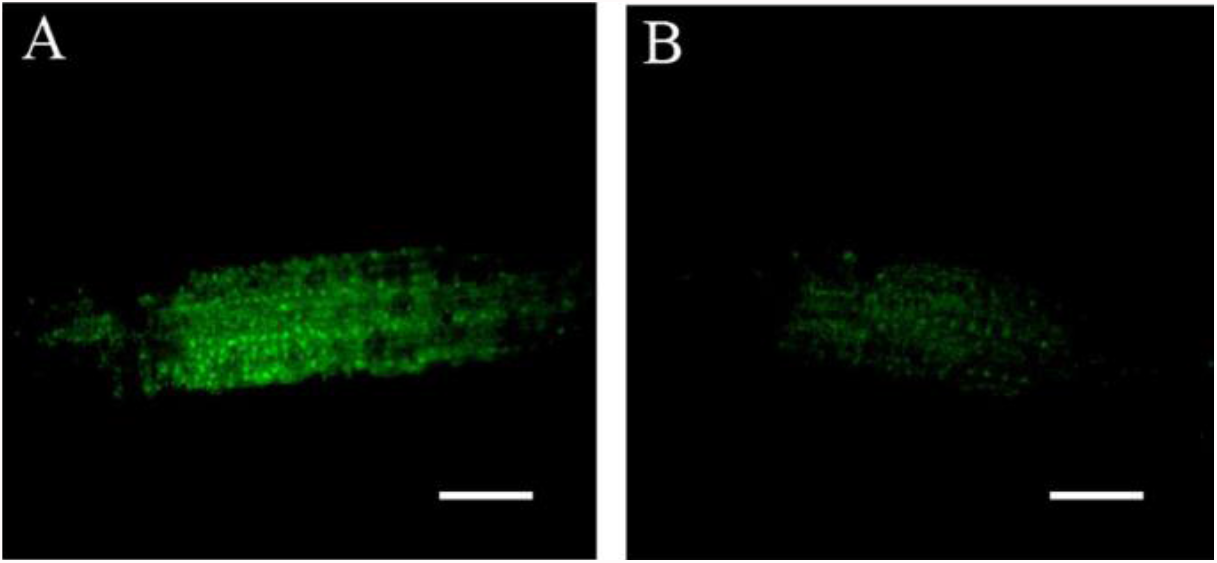
Auxin prevents the NaCl-induced increase of ABI4. Ten-day old seedlings were treated with 0.5 X MS 0.5% sucrose, and 0.3 M NaCl without (A) or with (B) 20 μM IAA. Roots were examined 2.5 h later. Scale bar = 100 μm.

### Steady-state levels of ABI4-eGFP are controlled by *de-novo* translation and degradation by the 26S proteasome

We have used the protein synthesis inhibitor cycloheximide (CHX) and the proteasome inhibitor MG132 to further characterize the transient accumulation of ABI4-eGFP following exposure to NaCl. As expected, CHX prevented the NaCl-dependent accumulation of ABI4-eGFP protein (Figure 8A), suggesting that exposure to NaCl enhances *de novo* translation of ABI4-eGFP. Treatment with a mix of NaCl and MG132 resulted in increased stabilization of ABI4-eGFP, and a high signal could be detected even 6 h after the co-application of NaCl and MG132 (Figure 8B), but not in the roots of plants treated with NaCl alone.

**Figure 8.**
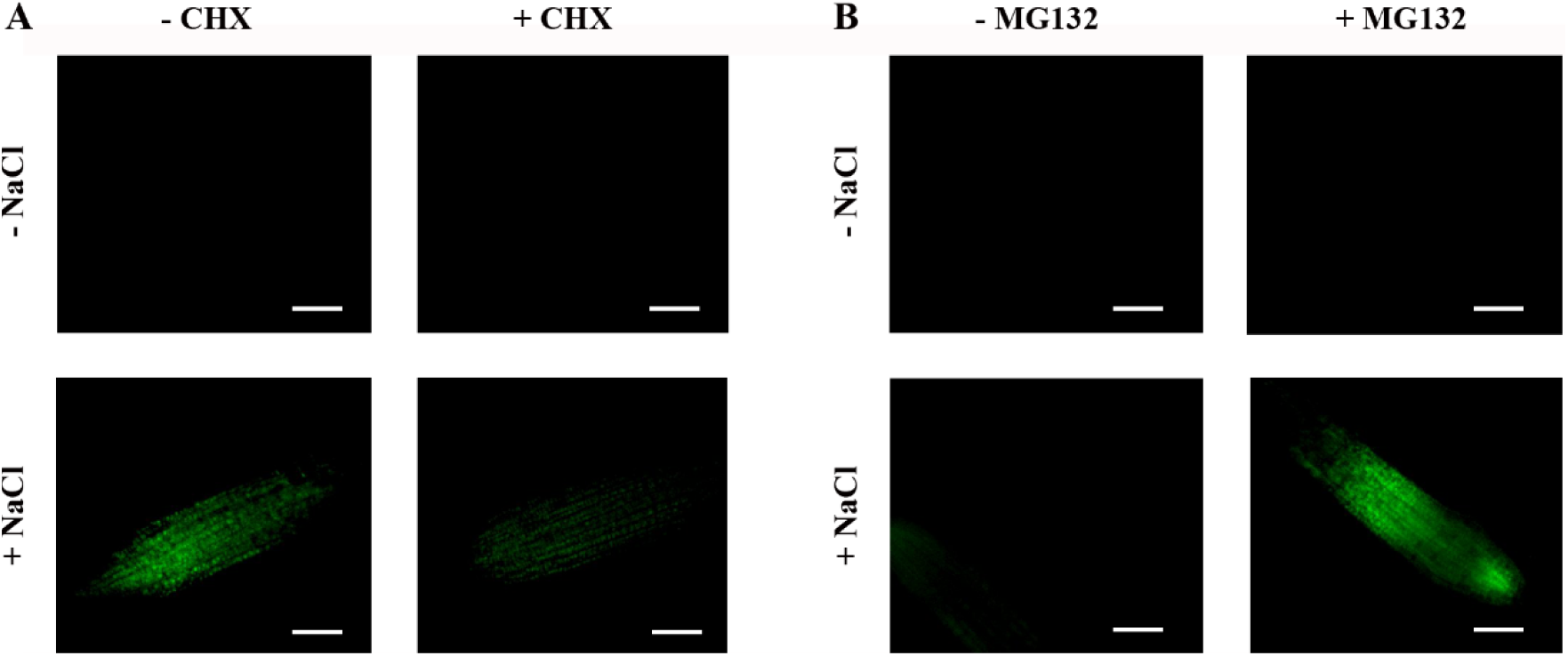
The NaCl-dependent transient increase in the ABI4-eGFP levels is a result of *de-novo* protein synthesis and its degradation by the 26S proteasome. Ten day old *ABI4-eGFP*-expressing seedlings were incubated in the light for 2.5 h (A) or 6 h (B) on filter paper soaked with 0.5 x MS, 0.5% sucrose solution supplemented, as indicated, with 0.3 M NaCl, 20 μg/ml cycloheximide (CHX), or 20 μg/ml MG132. Root were then examined by fluorescence microscopy. Scale bar = 100 μm.

### The phosphorylation state of serine 114 affects the stability of the ABI4 protein

We recently showed that phosphorylation of serine 114 of ABI4 by MPK3 or MPK6 is essential for its biological activity (Eisner *et al*., 2021). Here we tested whether the phosphorylation state of S114 of ABI4 also affects its stability, WT Arabidopsis plants were transformed with *35S::HA-FLAG-ABI4-eGFP* constructs encoding the ABI4 (S114A), phosphorylation null mutant or ABI4 (S114E), phosphomimetic mutated proteins. Ten day old NaCl treated seedlings were examined by fluorescent microscopy. Roots expressing WT (114S) ABI4-eGFP showed very low levels of fluorescence (Figure 9A), as in Figure 3. Fluorescence levels in roots of plants transformed with the ABI4-eGFP (S114A) phosphorylation-null mutant (Figure 9B) were similar to those of the WT ABI4-eGFP protein. In contrast, the S114E phosphomimetic mutation stabilized the ABI4-eGFP protein and high levels were observed even at 6 h following NaCl exposure (Figure 9C), indicating that the phosphorylation of serine 114 by MAPKs stabilizes ABI4, the active form of this transcription factor.

**Figure 9.**
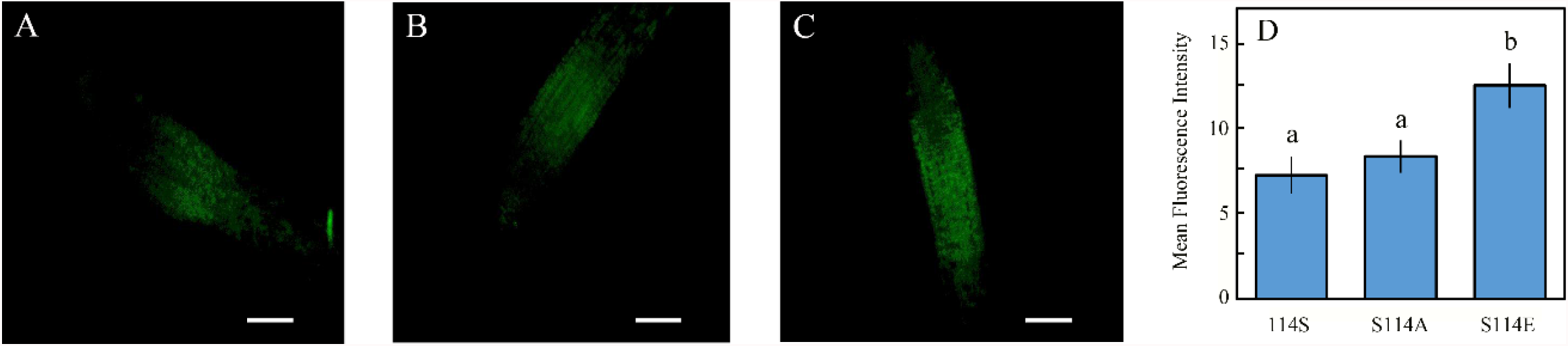
Phosphorylation of S114 stabilize ABI4 protein. Ten-days old ABI4-eGFP-expressing seedlings expressing: (A), WT ABI4-eGFP; (B), the phosphorylation null (S114A) mutant; or (C), the phosphomimetic (S114E) mutant, were incubated for 6 h with 0.5 x MS, 0.5% sucrose and 0.3 mM NaCl. The roots were examined by fluorescence microscopy. Scale bar = 100 μm. (D) The fluorescent signals of 70 plants were quantified. Data shown average ± SE. Bars with different letters represent statistically different values by Tukey’s HSD post-hoc test (P < 0.05).

## DISCUSSION

ABI4 is a master transcription regulator acting as both activator and repressor in the regulation of developmental processes such as seed development, germination, and root development, response to stress and hormones, disease resistance, and lipid metabolism (Wind *et al*., 2013; Chandrasekaran *et al*., 2020). *ABI4* is evolutionarily conserved and is a single gene in Arabidopsis and in most plant genomes that encode ABI4 (Wind *et al*., 2013; Xie *et al*., 2019), suggesting that its biological role is non-redundant. As a result, *abi4* mutants display pronounced phenotypes such as insensitivity to ABA inhibition of seed germination, reduced sensitivity to high glucose and salt (Finkelstein, 1993; Arenas-Huertero *et al*., 2000; Quesada *et al*., 2000; Shkolnik-Inbar and Bar-Zvi, 2010).

### *ABI4* is a lowly expressed and highly regulated gene

As expected for a key regulator, ABI4 levels and activity are tightly regulated. Maximal levels of *ABI4* transcripts are detected in developed seeds and at early germination stages with very low levels being present at other developmental stages (Söderman *et al*., 2000; Nakabayashi *et al*., 2005; Penfield *et al*., 2006; Shkolnik-Inbar and Bar-Zvi, 2011) where it is expressed in the phloem and parenchyma of the roots (Shkolnik-Inbar and Bar-Zvi, 2010; Shkolnik-Inbar and Bar-Zvi, 2011). ABI4 expression is regulated by plant hormones: enhanced by ABA (Söderman *et al*., 2000), cytokinin (Huang *et al*., 2017) and reduced by auxin (Shkolnik-Inbar and Bar-Zvi, 2010). It is also enhanced in response to high glucose (Arroyo *et al*., 2003), and osmotic (Arroyo *et al*., 2003) and salt (Shkolnik-Inbar *et al*., 2013) stresses. Arabidopsis *ABI4* is encoded by an intronless gene. Interestingly, intronless genes characterize highly regulated TFs in both plants and animals (Aviña-Padilla *et al*., 2021; Liu *et al*., 2021). Moreover, intronless genes are differentially expressed in response to drought and salt treatment (Liu *et al*., 2021).

### ABI4 is a post-transcriptionally regulated low-level protein

Since the transcription of *ABI4* is highly regulated, to study the post-transcriptional regulation of ABI4, we expressed *HA-FLAG-ABI4-eGFP* driven by the constitutive highly expressed CaMV 35S promoter (35S). eGFP is a GFP variant that is 35 times brighter than the original GFP (Zhang *et al*., 1996) thus allowing the detection of lower concentrations of tagged proteins than with the previously used ABI4-GFP (Finkelstein *et al*., 2011). This construct could complement the phenotype of the *abi4*-1 mutant, indicating that tagging ABI4 at its N- and C-termini does not eliminate its biological activity (Figure 1). We could not detect recombinant ABI4 protein in ten-day-old seedlings grown on plates under control conditions (Figure 2). In contrast, a high fluorescence signal is observed in imbibed embryos, confirming the performance of the construct (Figure 2). Although we used the highly active constitutive viral 35S CaMV promoter to express ABI4-eGFP, the resulting transgenic plants did not show any significant fluorescence of the eGFP tag (Figure 2). GFP-ABI4 could not be detected in Arabidopsis plants transformed with 35S::GFP-ABI4 (Finkelstein *et al*., 2011). Interestingly, GUS activity staining could identify the expressing ABI4-GUS recombinant protein driven by the same promoter. In contrast, ABI4-GFP fluorescence was detected in Arabidopsis protoplasts transfected with a *35S::ABI4-GFP* construct (Gregorio *et al*., 2014). This discrepancy may be explained by protoplasts being under stress caused by enzymatic digestion of the cell wall (Birnbaum *et al*., 2003).

### ABI4 is stabilized by external signals

Although the 35S promoter is active in most plant tissues, ABI4-eGFP is expressed primarily in the roots following stress (Figure 3), confirming the observations by Finkelstein et al who expressed ABI4-GUS fusion protein (Finkelstein *et al*., 2011). ABI4-eGFP was observed mainly in the vascular system of the roots following ABA and glucose treatments (Figure 6). Interestingly, ABI4-eGFP accumulated in the cells in which the endogenous ABI4 promoter is active (Shkolnik-Inbar and Bar-Zvi, 2010). Although NaCl treatment resulted in the accumulation of ABI4-eGFP throughout the roots, it was targeted to the nuclei only in the vascular cells (Figure 5), suggesting that both the accumulation and subcellular localization of the ABI4 protein is regulated in a cell-specific manner. This is similar to the transcription factor ABI5 that also accumulated following NaCl and ABA treatments, although increased levels were observed only four days after exposure to 200 mM NaCl (Lopez-Molina *et al*., 2001). Furthermore, ABA stabilization of ABI5 was restricted to a narrow developmental window 2 days after germination (Lopez-Molina *et al*., 2001). The difference in kinetics and responsive window suggests that although ABI5 and ABI4 proteins are stabilized by similar agents (ABA and NaCl), each protein has different domains (Finkelstein *et al*., 1998; Finkelstein and Lynch, 2000) and as such they are likely to be stabilized through different mechanisms. Transient expression of stress-induced genes has been reported for many genes, where the steady-state levels of mRNA peak at a given time after application of the stress agent, followed by a decrease. For example, mRNA levels of the stress-induced Arabidopsis transcription factors DREB1A DREB2A and rd29A are transiently induced following exposure to cold, drought, and salt stresses (Liu *et al*., 1998).

### Phosphorylation of S114 stabilizes ABI4

The phosphomimetic S114E form of ABI4 was more stable than the WT or the non-phosphorylated S114A mutant (Figure 9), suggesting that phosphorylation of S114 may decrease its ubiquitylation by a yet unidentified ubiquitin ligase. The S114 residue is included in the serine/threonine (S/T) region motif of ABI4 (Finkelstein *et al*., 1998). Several domains are proposed to contribute to the instability of ABI4: the PEST domain located at the N-terminus of ABI4 (amino-acids 22-40) enhances the degradation of ABI4 (Finkelstein *et al*., 2011; Gregorio *et al*., 2014). Furthermore, the N-terminal half of the ABI4 protein, including the PEST, APETALA2 (AP2), serine/threonine rich domain (S/T), glutamine-rich domain (Q), and the C-terminal half containing the Q and proline-rich (P) domains were shown to be highly unstable (Finkelstein *et al*., 2011). Degradation of the N-terminal half but not the C-terminal half of ABI4 was suppressed by the MG132 proteasome inhibitor, suggesting that although highly unstable, the C-terminal half of ABI4 may not be degraded by the proteasome (Finkelstein *et al*., 2011). The AP2-associated motif was also shown to destabilize ABI4 (Gregorio *et al*., 2014). Although the S/T rich region was included in the labile N-terminal half of ABI4 (Finkelstein *et al*., 2011), the instability of this region was mainly attributed to the PEST motif. Proteasomal degradation of ABI4 through the PEST motif is modulated by sugar levels (Gregorio *et al*., 2014).

Using the proteomic approach in human cell lines, Wu et al. (Wu *et al*., 2021) recently showed that phosphorylation delays the turnover of many proteins in growing cells. Moreover, the phosphomimetic mutated proteins catenin beta-1 (CTNNB1) S191D and the transcriptional receptor protein YY1 S118D were more stable than the WT proteins, and the phosphorylation-null in which the respective serine residues were mutated to alanine were destabilized (Wu *et al*., 2021). In addition, phosphoserine residues had a larger stabilization effect than phosphothreonine, and phosphotyrosine had only a marginal stabilization effect.

Phosphorylation of Type-A Response Regulator 5 (ARR5) by SnRK2s enhanced its stability (Huang *et al*., 2018). Furthermore, overexpressing WT ARR5 but not the non-phosphorylatable mutated-protein enhanced ABA hypersensitivity suggesting that the phosphorylated form of ARR5 is biologically active. ABA suppressed the degradation of ARR5 (Huang *et al*., 2018). Phosphorylation of the rate-limiting enzyme of ethylene biosynthesis, 1-aminocyclopropane-1-carboxylic acid synthase2 and 6 (ACS2 and ACS6) by MPK6 stabilizes the respective ACS proteins. Furthermore, the phosphomimetic ACS6 mutant was constitutively active, suggesting that phosphorylation of ACS6 by MPK6 is essential for its activity (Liu and Zhang, 2004). The RNA binding protein Tandem Zinc Finger 9 (TZF9) is destabilized by MAPK-mediated phosphorylation (Maldonado-Bonilla *et al*., 2014).

### MAPK regulates ABI4 both transcriptionally and post-transcriptionally

We show that phosphorylation of S114 stabilizes ABI4 (Figure 9). We recently demonstrated that MPK3, MPK4, and MPK6 phosphorylate S114 of ABI4 and that this phosphorylation is essential for the biological activity of ABI4 and the complementation of *abi4* mutant plants (Eisner *et al*., 2021). MPK3, MPK4, and MP6 are involved in the abiotic and biotic stress response (reviewed by (Bigeard and Hirt, 2018). Interestingly, treatments such as NaCl, ABA, and high glucose, which result in stabilization of ABI4 (Figures 3 & 6), also enhance the steady-state levels of the *ABI4* transcripts (Arroyo *et al*., 2003; Shkolnik-Inbar and Bar-Zvi, 2010; Shkolnik-Inbar *et al*., 2013). Here our results indicate that MAPK signaling affects both ABI4 transcription and protein stability.

The kinetics we found for the transient stabilization of ABI4 following salt treatment (Figure 3) resembles the described transient activation of MKK5 following exposure of Arabidopsis plants to high salt where increased activity of MKK5 is detected within 30 min of the treatment, reaching maximal activity at 2-4 h, and declining at 6 h after exposure to NaCl to nearly basal activity levels (Xing *et al*., 2015). MKK5 phosphorylates and activates several MPKs, including MPK3, MPK4, and MPK6. Therefore, the activity of these MPKs is also expected to be transient following salt treatment, resulting in a transient wave of phosphorylation of ABI4. MPK4 and MPK6 are rapidly activated by treatments such as high salt and osmotic stress but not by ABA treatment (Ichimura *et al*., 2000). ABA activates the transcription of many genes encoding components of the MAPK cascade (Menges *et al*., 2008), suggesting that the slow kinetics leading to accumulation of ABI4-eGFP following ABA treatment may result from slow de novo synthesis of the MAPKs rather than fast activation of preexisting latent enzymes.

MPK3, MPK4, and MPK6 also phosphorylate the transcription factors WRKY and MYB (Popescu *et al*., 2009; Sheikh *et al*., 2016). Several WRKY and MYB transcription factors may regulate ABI4 expression (Shang *et al*., 2010; Reeves *et al*., 2011; Feng *et al*., 2014; Ding *et al*., 2015; Lee and Seo, 2015; Huang *et al*., 2016; Chen *et al*., 2017; Ma *et al*., 2019; Guo *et al*., 2020; Li *et al*., 2021). In addition, as ABI4 also activates the transcription of its own gene (Bossi *et al*., 2009), its phosphorylation by these MAPKs also enhances its transcript levels.

In summary, our results show that phosphorylation of ABI4 by MAPK results in the stabilization of ABI4. Phosphorylation of S114 by MPKs may interfere with its binding to a yet unidentified E3 for proteasomal degradation. Alternatively, the catalytic efficiency of the E3 may be reduced towards phosphorylated ABI4. MAPK signaling also regulates ABI4 transcription. Thus, we suggest that regulation of both the *ABI4* transcript and ABI4 protein levels results in the tight regulation of the activity of this key transcription factor in the ABA signaling pathway.

## EXPERIMENTAL PROCEDURES

### Plant Material and Growth Conditions

*Arabidopsis thaliana* (Col) seeds of the indicated genotypes were surface sterilized, cold treated for 3 days, and plated in Petri dishes containing 0.5 X Murashige and Skoog medium (MS), 0.55% Plant Agar and 0.5% (w/v) sucrose as previously described (Shkolnik-Inbar and Bar-Zvi, 2010). Plates were incubated at 22-25 °C, 50% humidity under a circadian regime of 12 h light 12 h dark.

### Constructs and Plant Transformation

pGA-eGFP2 vector was constructed by replacing the sequences of the MCS and 35S::mGFP5 (9640-1038) in the pCAMBIA1302 vector (www.cambia.org) with 2 × 35S-MCS-eGFP DNA sequence (405-2332) from the pSAT4-eGFP-N1 plasmid by using Gibson assembly cloning (Gibson *et al*., 2009). The DNA sequence encoding HA_3_-FLAG_3_-ABI4 was isolated by digesting the pJIM19-ABI4 plasmid (Shkolnik-Inbar *et al*., 2013) with restriction enzymes *Nco*I and *Pst*I, and subcloning into the respective sites in pGA-eGFP2, to yield the pGA-HA3-FLAG3-ABI4 plasmid. To construct plasmids encoding ABI4 (S114A) and ABI4 (S114E) mutant proteins, their DNA sequences were amplified from the respective pRSET-ABI4 plasmid (Eisner *et al*., 2021) using gene specific primers flanked by the *Sal*I restriction sites, and digesting the amplified sequences with *Sal*I. The DNA sequence encoding WT-ABI4 was removed from the pGA-HA3-FLAG3-ABI4 plasmid by digestion with *Sal*I, followed by subcloning of the DNA sequences encoding mutated ABI4 protein. Primers used for the construction of plasmids are shown in Table S1.

The resulting plasmids were verified by PCR and DNA sequencing and were introduced into *Agrobacterium tumefaciens* strain GV3101. The transformed bacteria were used to transform WT Col or *abi4-1* Arabidopsis plants by the floral dip method (Clough and Bent, 1998). Transgenic plants were selected on plates containing hygromycin, and transferred to pot. Plant were grown at 22-25 °C, 50% humidity with 16 h light 8 h dark. Homozygous T2 and T3 generation plants were used in this study.

### Germination assay

Sterilized cold treated seed were plated on agar-solidified 0.5 × MS, 0.5% sucrose medium supplemented with the indicated concentrations of the phytohormone ABA. Germination was scored 7-days later.

### Plant treatment

For the different treatments, 10 days old seedlings were transferred to Petri dishes containing Whatman No.1 filter papers soaked with 0.5 × MS medium and 0.5% (w/v) sucrose supplemented with the indicated stress agent, plant hormone or inhibitors. Plants were incubated at room temperature in the light for the indicated times.

### Microscopy

The indicated tissues were examined using a fluorescent microscope (ECLIPSE Ci-L; Nikon) using filters set for GFP. All pictures taken in each experimental repeat were taken using the same microscope and camera setup and exposure times. Each experiment was repeated at least 3 times using at least 4 independent lines of the transgenic plants. Subcellular localization images were taken by a 3I Marianas spinning disc confocal microscope (Axio-observer 7 inverter; Zeiss) equipped with a Yokogawa W1 module and prime 95B sCMOS camera.

### Fluorescent quantification

Fluorescent signals were quantified using ImageJ software (Rasband, 1997-2015), with the black background set as zero for measurement of the fluorescent intensity of the image.

### Embryo excision

Arabidopsis seeds imbibed for 24 h in water at room temperature, were pressed gently between two microscope slides. Embryos released from seed coats were collected and rinsed briefly in water.

### Total protein extract, SDS-PAGE and western blot analysis

Ten-day-old seedlings were harvested into a 1.5 ml microcentrifuge tube, and their fresh weight was determined. 2:1 (v/w) 4 × SDS-PAGE sample buffer (Laemmli, 1970) was added, and the seedlings were homogenized with a microcentrifuge pestle. To ensure efficient solubilization of plant proteins, homogenates were passed through 2 cycles of freezing in liquid nitrogen and boiling for 5 min, and then heated for 5 min. Tubes were centrifuged for 10 min at 12,000 × g at room temperature, and supernatant samples were resolved by SDS PAGE. Proteins were electroblotted onto nitrocellulose membranes. ABI4-eGFP and β-actin were detected using the primary antibodies anti-GFP (Abcam, ab1218) and anti-β-actin (Sigma, A4700), respectively, and secondary peroxidase-coupled anti-mouse IgG antibody (Sera Care 5450-0011). Membranes were incubated in reaction mix prepared from the highly sensitive SuperSignal West Dura extended substrate kit (Thermo scientific, 34075), and chemiluminescent signals were recorded using ImageQuant RT ECL Imager (GE Healthcare).

### Quantitative RT-PCR analysis

Total RNA was isolated from roots using a ZR Plant RNA MiniPrep kit (Zymo research). The RNA concentration was estimated spectrally (Nano Drop ND-1000; Nano Drop Technologies). cDNA was synthesized using the qScript cDNA synthesis kit (Quanta). The reaction mixture contained 700 ng of total RNA and random primers. Primer design and RT-qPCR assays for determining relative steady state transcript levels were as previously described (Shkolnik-Inbar *et al*., 2013). Primers are described in Table S1.

## ACCESSION NUMBERS

ABI4, At2G40220

## ACKNOWLEDGEMENTS

We thank Guy Adler for constructing the pGA-eGFP vector. This work was supported by grant No. 2011097 from the US Israel Binational Science Foundation (BSF) (to Dudy Bar-Zvi).

Dudy Bar-Zvi is the incumbent of The Israel and Bernard Nichunsky Chair in Desert Agriculture, Ben-Gurion University of the Negev.

## SHORT LEGENDS FOR SUPPORTING INFORMATION

**Figure S1**. eGFP expression in roots of salt-treated *35s::eGFP* plants. Ten days old transgenic plants overexpressing *35S::eGFP* incubated for the indicated times with 0.5 × MS, 0.5% sucrose, without NaCl (A) or with 0.3 M NaCl, for 2.5 h (B), 4 h (C), 6 h (D). Roots were examined by fluorescence microscopy. Scale bar = 100 μm.

**Table S1**. Primers used in this study.

