## Supplementary material for "The ABA INSENSITIVE (ABI) 4 transcription factor is stabilized by stress, ABA and phosphorylation": Figure S1

**Figure S1.** eGFP expression in roots of salt-treated *35s::eGFP* plants. Ten days old transgenic plants overexpressing *35S::eGFP* incubated for the indicated times with 0.5 x MS, 0.5% sucrose, without NaCl (A) or with 0.3 M NaCl, for 2.5 h (B), 4 h (C), 6 h (D). Roots were examined by fluorescence microscopy. Scale bar = 100 μm.

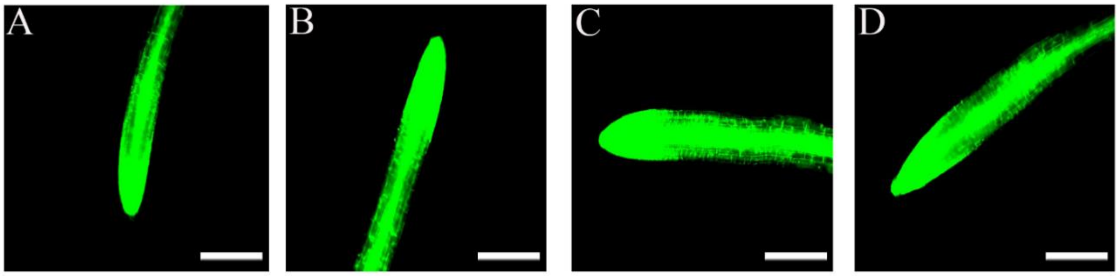
