## Supplementary material for "The ABA INSENSITIVE (ABI) 4 transcription factor is stabilized by stress, ABA and phosphorylation": Table S1

**Table S1**. Primers used in this study

| No | Primer set | Forward primer | Reverse primer | Used for |
|---|---|---|---|---|
| 1 | pCAMBIA 1302 | CTAATTCCTAAAACCAAAATCCAGTGACAATT AAACTATCAGTGTTTGACAGGATA | CATGTTGACCGGTTAGGGATAACAGTGCCTAATG AGTGAGCTAACTCAC | Construction of the pGA-eGFP plasmid |
| 2 | pSAT4 | GTGAGTTAGCTCACTCATTAGGCACTGTTATC CCTAACCGGTCAACATG | TATCCTGTCAAACACTGATAGTTTAATTGTCACTG GATTTTGGTTTTAGGAATTAG | Construction of the pGA-eGFP plasmid |
| 3 | ABI4 SalI | GTCGACCTCGAGATGGACCCTTTAGCTTCCC | GTCGACTGCAGATAGAATTCCCCCAAGATGGGAT | Amplification of sequences encoded mutated *ABI4* |
| 4 | TAP | ATGTACCCATACGATGTTCCTGAC | AAGCTTGATATCAGCGTAATCTGGA | RT-qPCR of *ABI4-eGFP* |
| 5 | 18S | AAGCAAGCCTACGCTCTGGA | AGGCCAACACAATAGGATCGA | RT-qPCR of 18S rRNA |
